# Suppressive effects of oroxylin A on intracellular proliferation of *Toxoplasma gondii* via host cell ERK phosphorylation inhibition

**DOI:** 10.1101/2024.03.17.585380

**Authors:** Ziyue Z Zhang, Kazumi Norose, Noriko Shinjyo, Xiaoxia X Lin, Akiko Suganami, Yutaka Tamura, Hirokazu Sakamoto, Kenji Hikosaka

## Abstract

*Toxoplasma gondii* poses a significant threat to immunocompromised patients, resulting in high mortality rates. Considering the side effects of anti-*Toxoplasma* drugs, we focused on a potential candidate, oroxylin A (OA), a common component extracted from *Astragalus membranaceus* and *Scutellaria baicalensis* which suppressed the growth of *T. gondii in vitro* and *in vivo.* Our result demonstrated that OA suppressed *T. gondii* intracellular proliferation and downregulated phosphorylation of ERK1/2 of *T. gondii*-infected host cells very similar to the MEK1 specific inhibitor PD98059. Our *in silico* analysis showed that OA interacts sufficiently with the mammalian MEK1 region where established MEK1 inhibitors like PD98059 and trametinib bind. Moreover, OA improved the survival rate in *T. gondii*-infected mice. This study proposes that host MEK1 is a novel potential target for anti-*Toxoplasma* drugs with a new mechanism of action.

---

*Toxoplasma gondii*, a protozoan parasite known for its remarkable success as an intracellular parasite, infects approximately one-third of the world’s population^1^ and is reported to infect nearly all warm-blood animals. In humans, toxoplasmosis caused by this parasite typically remains asymptomatic in patients with a typical immune system. However, in cases involving immunocompromised patients, such as those with human immunodeficiency virus infection, organ transplantation, or pregnancy, *T. gondii* becomes activated within the host’s organs, leading to severe diseases like toxoplasmic encephalitis, ocular toxoplasmosis, and *Toxoplasma* pneumonia^2–4^. In general, pyrimethamine (PYR) and sulfadiazine are clinically used to treat acute toxoplasmosis. However, their severe side effects and the emergence of drug resistance pose serious issues that require continued treatment^5–7^. Recently, considerable attention has been paid to developing novel drugs that offer reduced side effects for combating toxoplasmosis. Hence, it becomes imperative to identify various compounds capable of inhibiting the growth of *T. gondii* via diverse mechanisms of action.

In the last decade, Traditional Chinese medicine (TCM) has gained increasing recognition as an alternative treatment for infectious diseases due to its fewer side effects^8,9^. Artemisinin, extracted from *Artemisia annua*, has found widespread use in combating malaria^10^. Similarly, *Astragalus membranaceus* (Am) and *Scutellaria baicalensis* (Sb) have exhibited effectiveness in suppressing the proliferation of *T. gondii* proliferation *in vivo*^11^ and *in vitro*^12^. These East Asian herbal medicines (e.g., TCM) have been trusted for their safety and are used to treat various inflammatory and infectious diseases^12,13^. However, the properties of herbal products can considerably differ based on their sources, making quality control and evaluation of efficacy based on active constituents essential. Therefore, the identification of compounds within these plants suppresses *T. gondii* proliferation, and elucidation of their mechanisms of action is paramount in developing novel drugs against toxoplasmosis. Searching for potential active components within Am^14^ and Sb^15^, we identified oroxylin A (OA) as a common constituent.

OA, a flavonoid, demonstrates a broad spectrum of bioactivities showcasing its potential because of its anti-tumor, anti-virus, anti-protozoan parasite, anti-inflammation, anti-oxidation and anti-allergy activities in addition to organ protection^16,17^. Considering its multi-bioactivities, OA holds considerable potency for clinical applications^17^. In addition, our focus on OA led us to delve into the flavone backbone structure. Certain flavonoids, including OA, can inhibit the MAPK signaling pathway^18^. This pathway plays a critical role in governing cellular processes such as cell proliferation, differentiation, stress response, and apoptosis^19,20^. Interestingly, *T. gondii* infection triggers the activation of the host-cell MAPK pathway, and studies indicate that inhibiting this pathway suppresses parasite proliferation^21^.

In this study, we hypothesized that OA could suppress *T. gondii* proliferation by inhibiting the host-cell MAPK signaling pathway. To investigate this, we assessed the effects of OA on *T. gondii in vitro* and *in vivo*, along with monitoring alterations in the host-cell MAPK signaling pathway during parasite infection. Furthermore, utilizing *in silico* docking simulation and *in vitro* assays, we inferred the target molecule and mechanism through which OA operates. Our findings provide a novel strategy for the development of novel drugs combating toxoplasmosis.

## RESULTS

### OA suppresses *Toxoplasma* proliferation *in vitro*

Initially, we evaluated the effect of OA on the proliferation of host cells (Vero cells) using the sulforhodamine B (SRB) assay^22, 23^. At concentrations of 50 µM and below, OA did not demonstrate any significant effect (Figure 1). Next, we evaluated the effect of OA on cytotoxicity induced by *T. gondii* proliferation using a monolayer disruption assay^24^, with PYR as the control. The results depicted in Figure 2A and 2B revealed a substantial decrease in the percentage of bottom coverage due to *T. gondii* infection. However, OA exhibited a dose-dependent increase in this coverage, indicating that OA suppressed *T. gondii* proliferation. At a concentration of 50 µM, OA displayed a comparable effect to that of 2 µM PYR. These results indicate that OA suppresses the proliferation of *T. gondii*.

**Figure 1.**
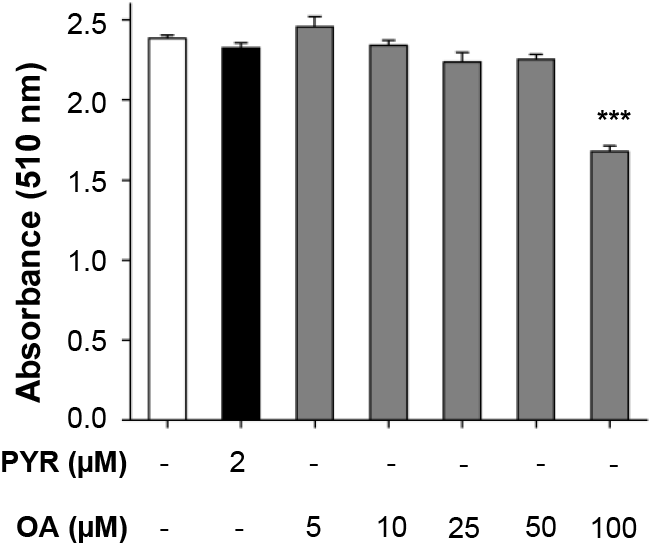
OA did not exhibit cytotoxicity towards host cells at concentrations of 50 µM and below. Evaluation of OA toxicity to host cells (Vero cells) by SRB assay. The white and black bars indicate the results of controls: no compound supplementation (DMSO-treated group) and PYR supplementation, respectively. The gray bars indicate the results of various OA concentration supplementations. The experiments were conducted three times independently and in triplicates. Results are depicted as mean ± SEM. ***, P < 0.001 vs. the DMSO-treated group.

**Figure 2.**
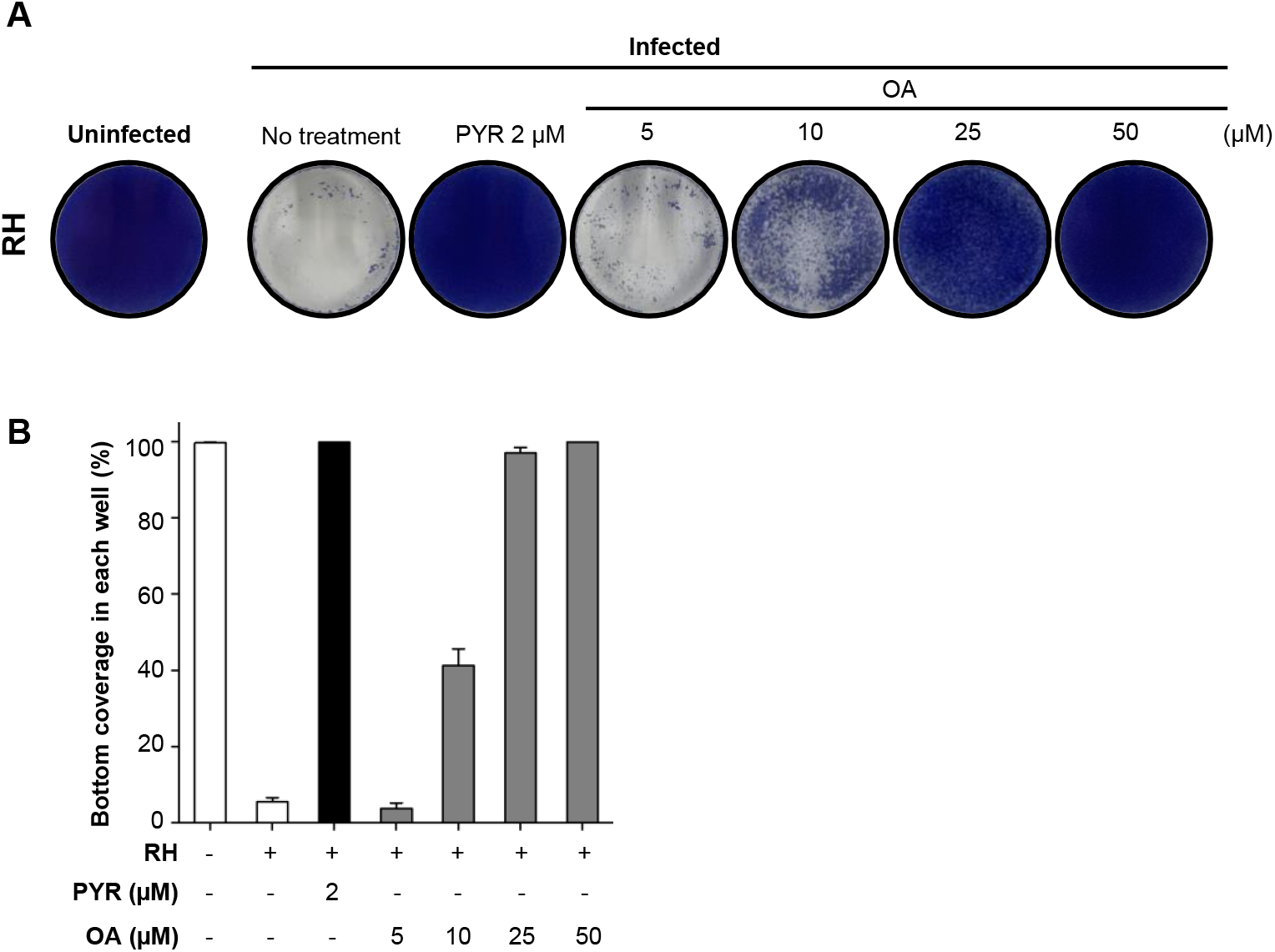
OA suppresses *T. gondii* proliferation. (**A**) Images of the monolayer disruption assay. Representative images for each condition are shown. The experiments were independently repeated twice and triplicated within each. **(B)** Comparison of the percentages of the bottom coverages for each condition. The white and black bars indicate the results of controls: no compound supplementation (DMSO-treated group) without and with *T. gondii* and PYR supplementation, respectively. The gray bars indicate the results of various OA concentration supplementations. The experiments were conducted three times independently and in triplicates. Data are presented as mean ± SD.

### OA suppresses the intracellular proliferation of *Toxoplasma*

To evaluate the mechanism of OA effect on parasite proliferation, we employed the *T. gondii* RH-GFP strain, engineered to express GFP constitutively, enabling the quantitative assessment of parasite number and size through GFP signals. We confirmed that OA suppressed the percentage of GFP signal density (Figure 3A and 3B). Thus, we utilized the RH-GFP strain in subsequent experiments.

**Figure 3.**
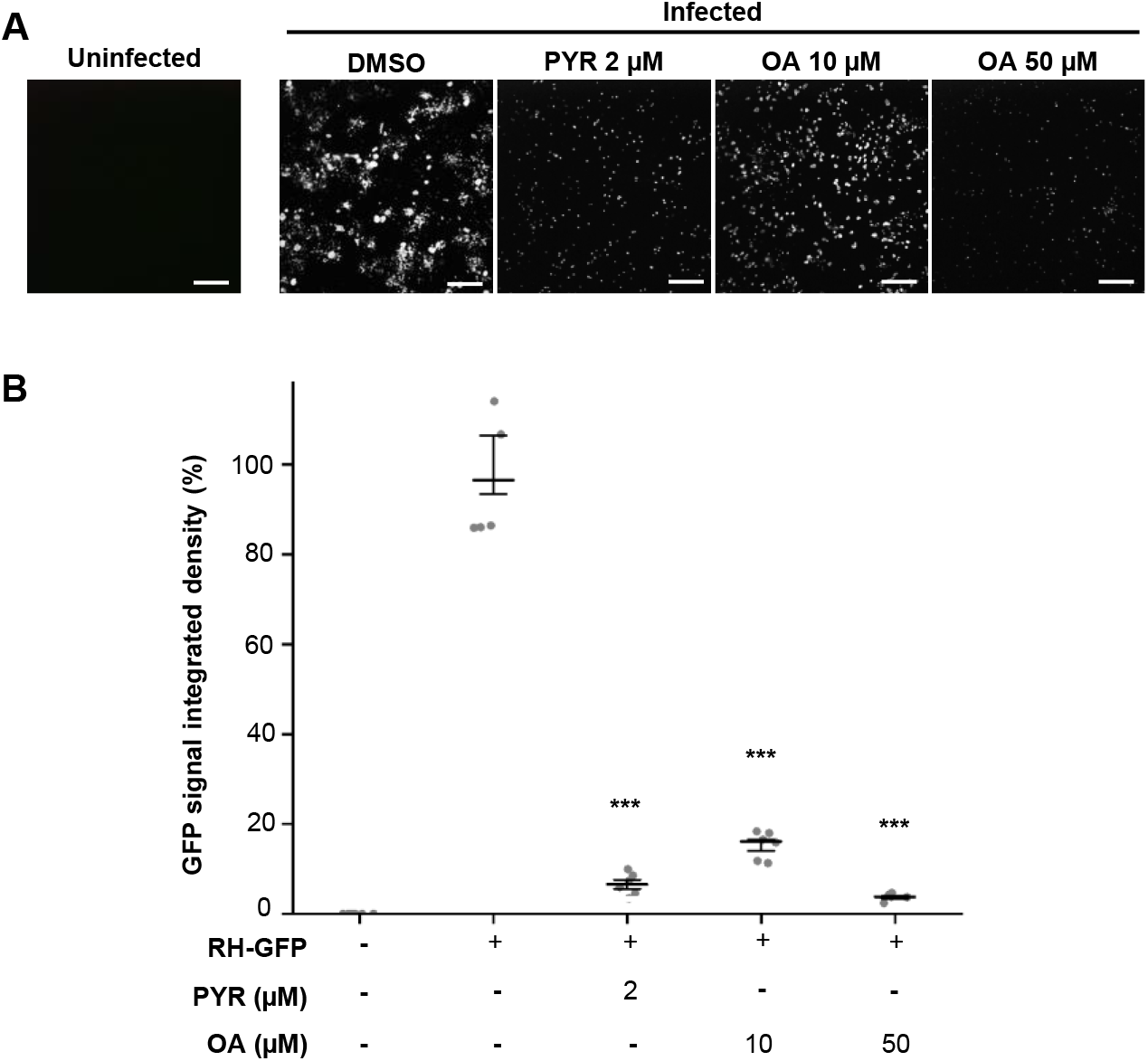
Assessing the effect of OA on *T. gondii* proliferation using the RH-GFP strain. (**A**) Representative images of GFP signals from uninfected and RH-GFP-infected host cells (Vero cells) at 48 h post-invasion. For the parasite-infected host cells, DMSO and PYR were used as negative and positive controls, respectively. Scale bar: 100 µm. **(B)** Comparison of the percentages of GFP signal-integrated density for each treatment. GFP signals were calculated from six random fields using ImageJ software. Data are presented as the median ± SD. ***, P < 0.001 vs. the infected DMSO-treated group.

To determine whether OA affects intracellular parasites, we measured the size of the parasitophorous vacuole (PV) within infected host cells. Results obtained at 30 h post-infection demonstrated a reduction in PV sizes due to the treatments (Figure 4A and 4B). In addition, it was noted that the PV size of parasites treated with 50 µM OA was significantly smaller compared to those treated with 10 µM OA (*P* < 0.001). These results suggest that OA suppresses the intracellular development of *T. gondii*.

**Figure 4.**
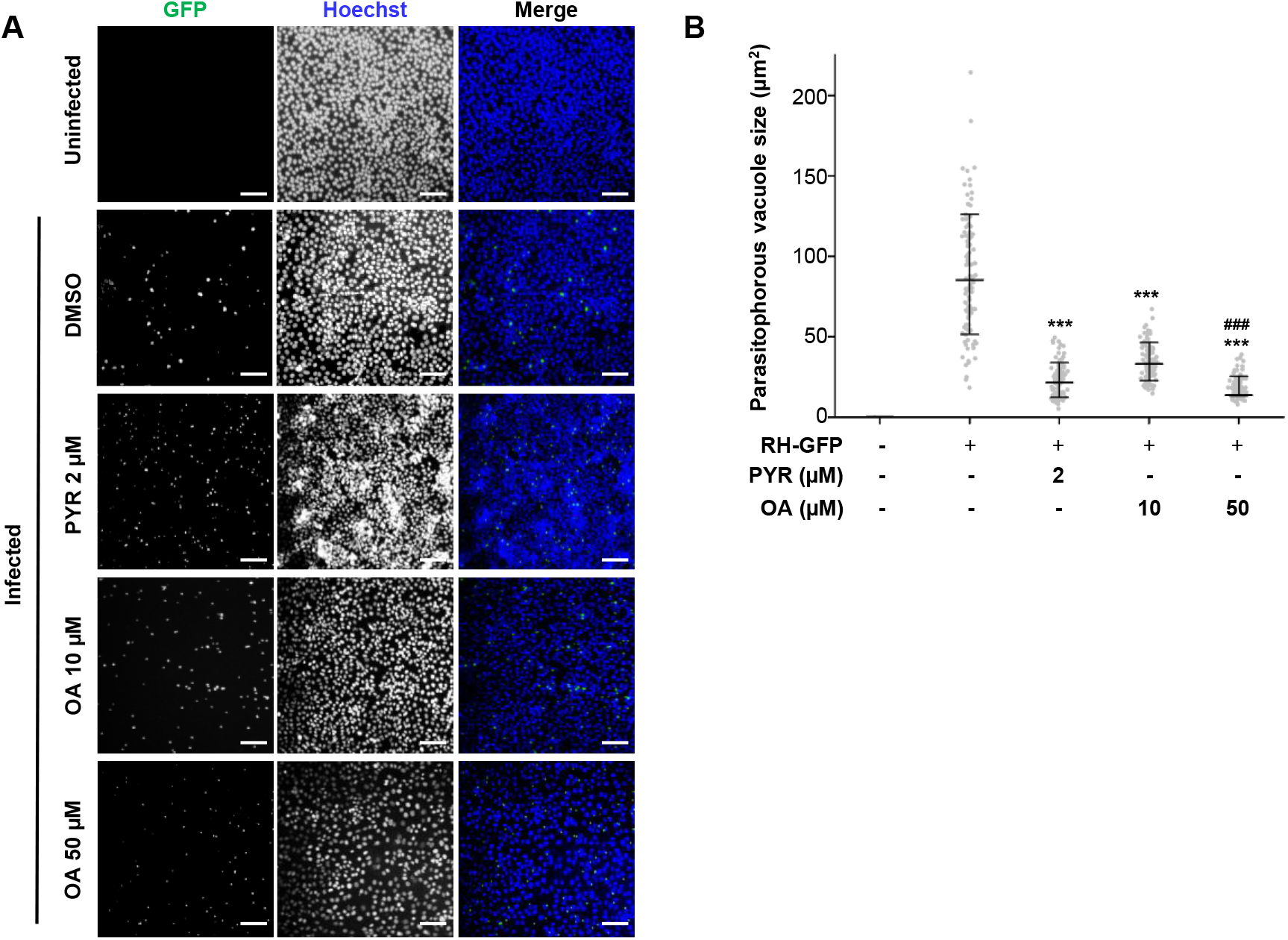
OA suppresses the intracellular development of *T. gondii*. (**A**) Representative images of GFP and Hoechst 33342 signals from uninfected and RH-GFP-infected host cells (Vero cells) at 36 h post-invasion. For the parasite-infected host cells, DMSO and PYR were used as negative and positive controls, respectively. The host cells were stained with Hoechst 33342. Scale bar: 100 µm. **(B)** The size of the area (µm^2^) of the parasitophorous vacuole (PV) across the indicated treatments. One hundred PVs in each condition were quantified using ImageJ software. Data are presented as the median ± SD. ***, P < 0.001 vs. the infected DMSO-treated group. ###, P < 0.001 vs. the infected 10 µM OA-treated group.

### OA does not affect extracellular *Toxoplasma* viability

Subsequently, we assessed the effect of OA on extracellular parasites. To assess this, we pretreated the extracellular parasites for 1 h with OA before invasion. At 48 h post-infection, there was no difference in the number of parasites between the pretreated and untreated conditions (Figure S1). Even upon extending the treatment duration to 4 h, encompassing 1 h before invasion and 3 h for invasion, no difference was observed between the conditions. These results suggest that OA does not influence the viability of extracellular parasites, including their motility and host-cell invasion activities.

### OA inhibits the host-cell MAPK signaling pathway induced by *Toxoplasma* infection

OA, a flavonoid characterized by a flavone backbone structure (Figure 5A), has demonstrated the ability to inhibit the MAPK pathway in osteoarthritis chondrocyte cells and non-small-cell lung cancer cells^19,20^. Furthermore, Han et al.^25^ indicates that *T. gondii* infection triggers the activation of the host-cell MAPK signaling pathway, and inhibiting this pathway results in decreased parasite proliferation in Vero cells. In addition, *T. gondii* lacks MAPK/ERK kinase (MEK) 1/2 homologs^26^. Therefore, we hypothesized that OA could target on the host-cell MAPK signaling pathway, but not on the parasite.

**Figure 5.**
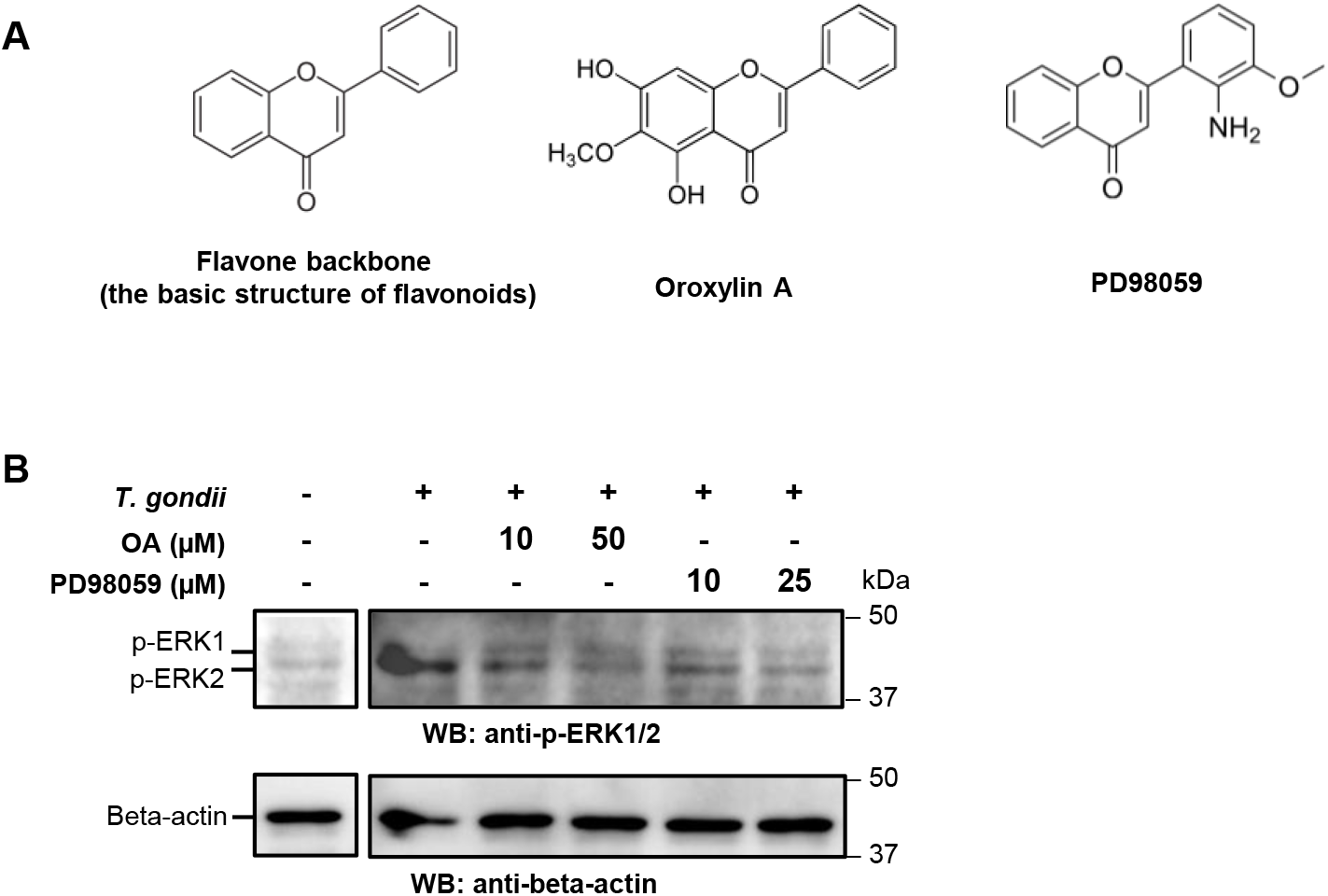
OA downregulates the host ERK1/2 phosphorylation in *T. gondii*-infected host cells. (**A**) The chemical structures of the flavone backbone, oroxylin A, and PD98059. **(B)** Comparison of the phosphorylation levels of host ERK1/2 in each treatment by western blotting. Phosphorylated ERK1/2 (p-ERK1/2) were detected using an anti-p-ERK1/2 antibody. Beta-actin was detected as a loading control.

To validate our hypothesis, we evaluated the activation of the MAPK signaling pathway in *T. gondii*-infected cells with and without OA treatment. We assessed the phosphorylation levels of extracellular signal-regulated kinase (ERK) 1 and ERK2 (Figure 5B), known as crucial endpoints within the MAPK pathway. We used PD98059 (Figure 5A), a specific inhibitor for MEK1^27^, as a positive control, which exhibited no cytotoxicity to the host cell at concentrations of 50 μM and below (Figure S2). We confirmed that host-cell ERK1/2 phosphorylation was increased upon *T. gondii* infection using Vero cells (Figure 5B) and human fibroblast cells (Figure S3) and the phosphorylation was decreased by PD98059 treatment (Figure 5B). We found that OA also inhibited infection-induced ERK1/2 phosphorylation (Figure 5B). These results demonstrated that OA counteracted the phosphorylation of host-cell ERK1/2 within the MAPK signaling pathway activated by *T. gondii* infection.

### A combination of OA and PD98059 does not have an additive or synergistic inhibitory effect on *Toxoplasma* proliferation

To confirm that the host MAPK pathway is significant for *T. gondi* growth, we examined the effect of PD98059 treatment on the parasite proliferation.We confirmed that PD98059 exhibited a dose-dependent inhibition of *T. gondii* proliferation in Vero cells (Figure 6A), as reported previously^19,25^.

**Figure 6.**
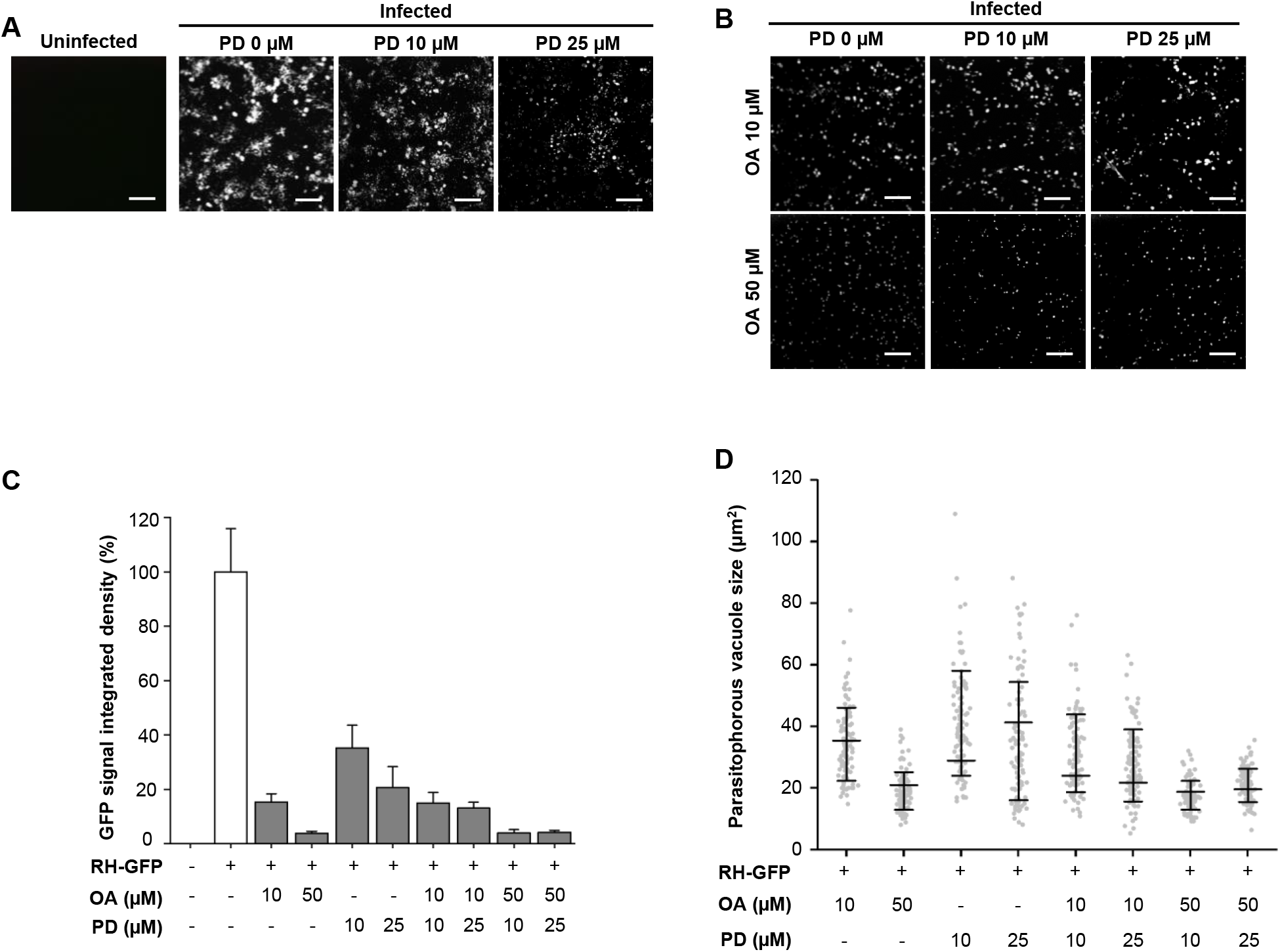
The combination treatment of OA and PD98059 does not exhibit an additional or synergistic inhibitory effect on *T. gondii* proliferation. **(A and B)** Representative images of GFP signals from uninfected and RH-GFP-infected host cells (Vero cells) treated with PD98059 (PD) **(A)** and single and combination treatments with OA and PD **(B)**. Scale bar: 100 µm. **(C)** The percentages of GFP signal-integrated density among the indicated treatments. GFP signals were calculated from six random fields using ImageJ software. Data are presented as mean ± SD. **(D)** The size of the area (µm^2^) of parasitophorous vacuole (PV) among the indicated treatments. One hundred PVs in each condition were quantified using ImageJ software. Data are presented as median ± SD. **(A–D)** These experiments were repeated three times independently and conducted in triplicate. PD, PD98059.

To elucidate which step of MAPK pathway is inhibited by OA, we investigated the impact of combining OA and PD98059 on the intracellular parasite proliferation. Our results indicated no significant additive or synergistic effect when combining 10 or 25 µM PD98059 with 10 and 50 µM OA, as observed in the GFP signal distribution (Figure 6B) and the % of GFP signal density (Figure 6C). Furthermore, the size of the PV housing the parasites did not exhibit any additive or synergistic upon adding 10 or 25 µM PD98059 to 10 and 50 µM OA (Figure 6D). These results suggest that OA and PD98059 share the same binding site within MEK1.

### *In silico* docking simulation supports that OA interacts with the host MEK1

To provide comprehensive insights into our *in vitro* experiments, we conducted an analysis to assess the physical and chemical affinity between OA and MEK1 using *in silico* docking simulation techniques. Known MEK1-specific inhibitors, PD98059^27^ and trametinib^28^, have been identified to bind to the allosteric site of MEK1. We compared the potential binding capacity of OA to MEK1 using PD98059 and trametinib as positive controls for MEK1 allosteric site-binding molecules. Subsequently, we constructed the three-dimensional structures of the trametinib/MEK1, PD98059/MEK1, and OA/MEK1 complexes closely evaluating the interaction between MEK1 and each ligand (Figure 7). Interaction energies between MEK1 and each ligand were trametinib (–111.55 kcal/mol) > PD98059 (– 60.79 kcal/mol, as a positive control) > OA (–54.21 kcal/mol), consistent with the results of the *in vitro* experiments (Figure 6). Moreover, our analysis identified specific amino acid residues within MEK1 commonly recognized by all ligands, D208, F209, V211, and S212 (Figure 7); these amino acids are conserved among mammals^29^. This results strongly support that OA binds to MEK1 in the same mode as the MEK1-specific inhibitors.

**Figure 7.**
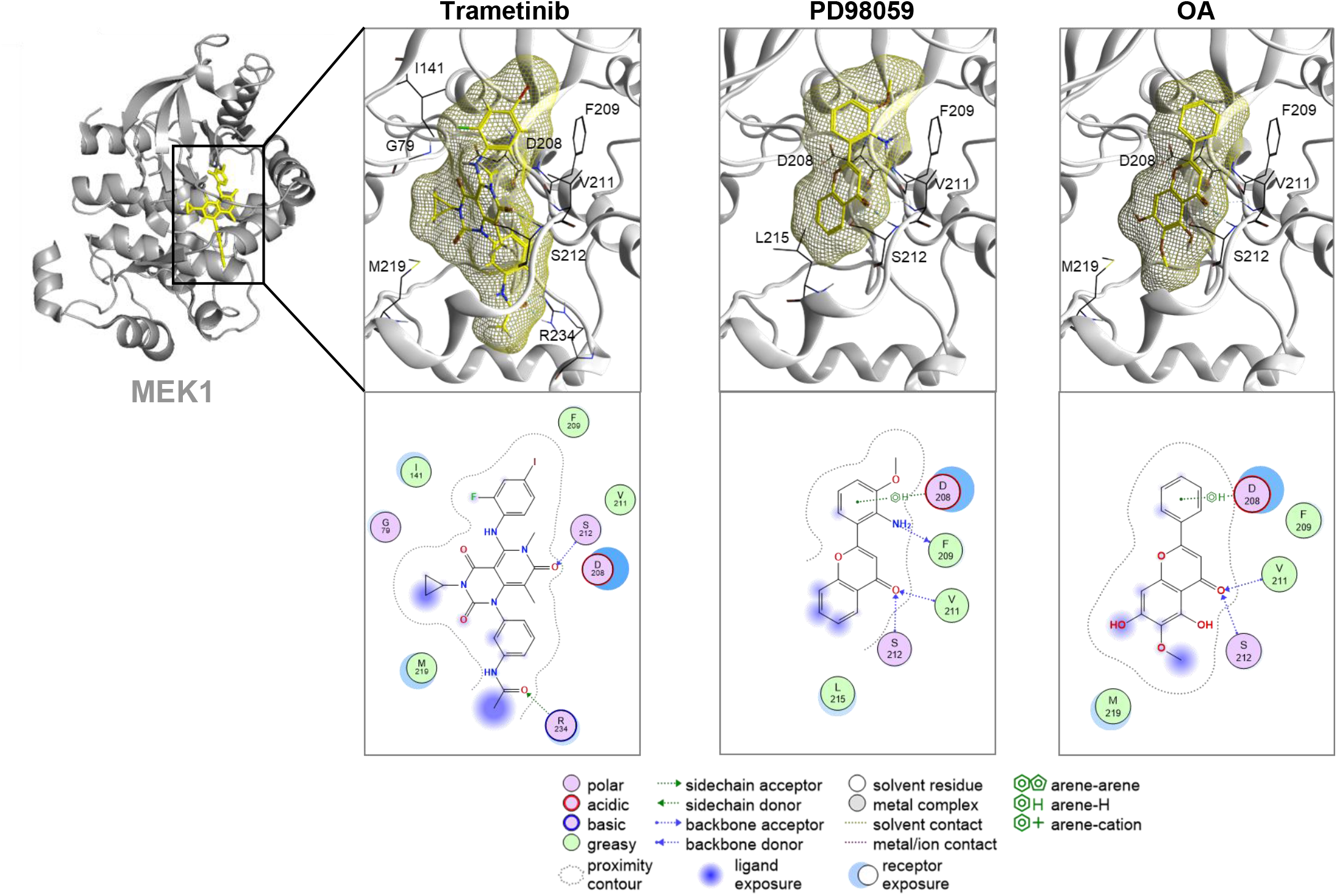
*In silico* docking simulation of ligand-bound complexes involving MEK1. The results of the docking simulations between the MEK1 (gray) allosteric site and indicated compounds (yellow). Trametinib and PD98059, known MEK1-specific inhibitors, are used as controls for the simulation. The three-dimensional (upper panel) and two-dimensional structures (lower panel) are shown. D208, F209, V211, and S212 in MEK1 are commonly involved in the binding to the compounds.

### OA treatment increases the survival rates of *Toxoplasma*-infected mice

Finally, we performed mouse infection experiments to assess the efficiency of OA in suppressing *T. gondii* growth and the safety of OA *in vivo*. We used a cyst-forming Fukaya strain (archetypal type II), closely resembling the clinical scenario observed in humans. We confirmed that OA suppressed the growth of this strain *in vitro* (Figure S4).

We monitored the survival rates of infected mice treated with or without OA, PYR, and sulfadiazine for 30 days (Figure 8A). In our experimental condition, all mice in the challenge control group (infected DMSO treated; n = 8) succumbed by 11 days post-infection (dpi.) (Figure 8B, black line). In contrast, all mice in the cure control group (infected P+S treated) survived until 30 dpi (Figure 8B, magenta line). On the other hand, around 40% of the mice in the infected OA-treated group (n = 8) survived at 11 dpi and they continued to survive until 30 dpi (Figure 8B, green line) (*P* < 0.06 vs the challenge control). This suggests that OA suppresses parasite growth also *in vivo*.

**Figure 8.**
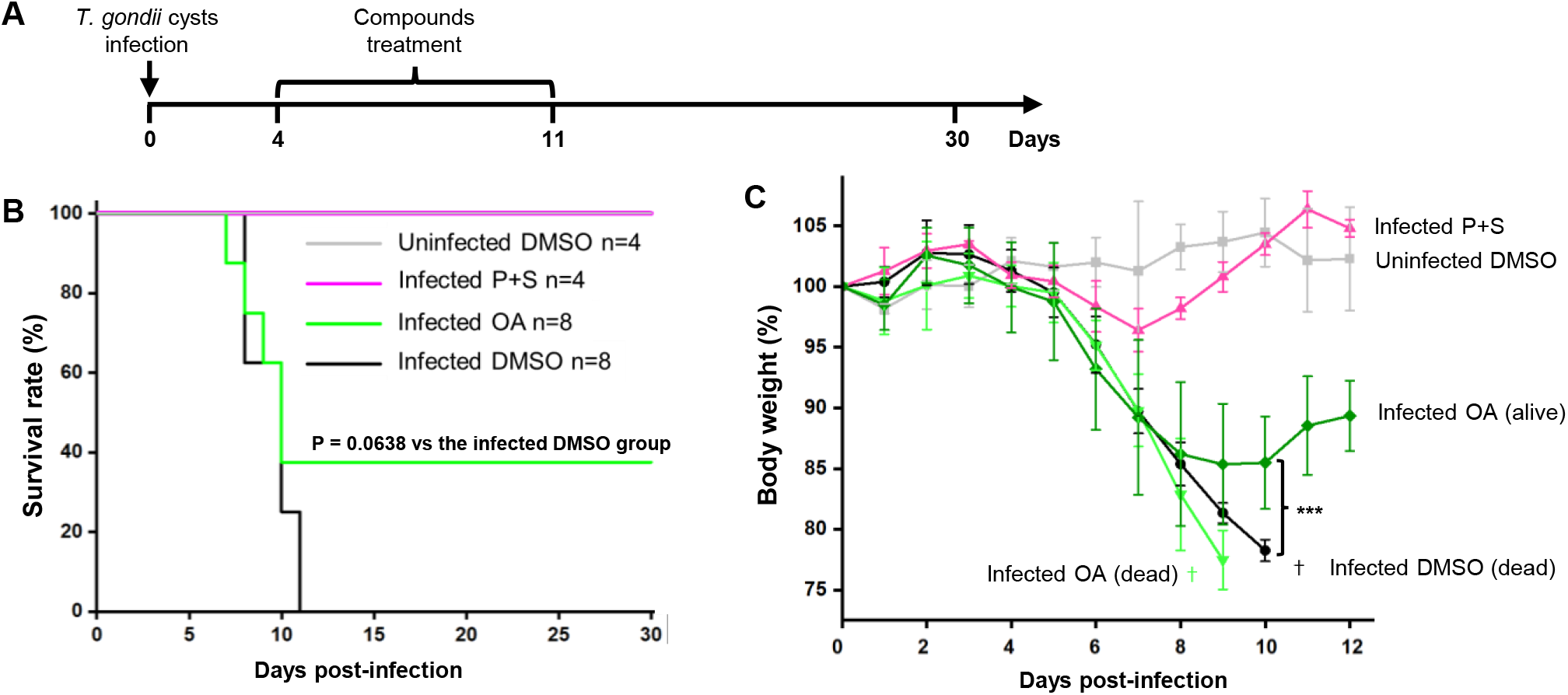
OA increased the survival rate of *T. gondii*-infected mice. **(A)** Experimental schedule of compound treatments. The *T. gondii* Fukaya strain was infected at day 0 and the compounds were treated during day 4 to 11. The mice were then monitored for survival until day 30. **(B)** Survival curve depicting uninfected DMSO (n = 4), infected DMSO (n = 8), infected P + S (n = 4), and infected OA (n = 8) treated mice for 30 days. P = 0.0638 between the infected DMSO-treated group and the infected OA-treated group. **(C)** Changes in body weight of mice among the indicated conditions. Data are presented as mean ± SD. ***, P < 0.001 vs. infected DMSO-treated group. DMSO, solvent; P + S, PYR, and sulfadiazine.

In addition, we monitored the effect of drug treatments on weight change (Figure 8C). In the infected DMSO-treated group, weight loss continued from day 4 and resulted in death.

In contrast, the infected P+S-treated group stopped losing weight on day 7 and began regaining weight from day 8. This suggests that P+S treatment eliminated *T. gondii* by day 7. In the OA-treated survival group, weight loss stopped on day 8 and recovered weight from day 10, and they continued to survive until 30 dpi. These suggest that *T. gondii* was eliminated by about day 8 and that there were no fatal side effects of OA treatment on survival, although the effects of OA were slower than those of P+S. These results indicate that OA suppresses parasite growth *in vivo* with low side effects.

## DISCUSSION

Developing new therapeutic drugs to combat toxoplasmosis requires identifying a broader array of safe and efficient hit and lead compounds^31^. The traditional East Asian traditional herbal medicines *Astragalus membranaceus* (Am) and *Scutellaria baicalensis* (Sb) inhibit *T. gondii* proliferation *in vitro* and *in vivo*^11,12^. This study focused on oroxylin A (OA), a shared constituent found in Am and Sb. We evaluated its potential as a new medicine against toxoplasmosis, where we demonstrated that OA suppressed *Toxoplasma* intracellular proliferation by inhibiting the host-cell MAPK signaling pathway (Figure 9). Our findings demonstrated that OA is one of the active components in Am and Sb for their anti-*T. gondii* activity. Furthermore, our study proposes that host MEK1 is a novel potential target for anti-*Toxoplasma* drugs.

**Figure 9.**
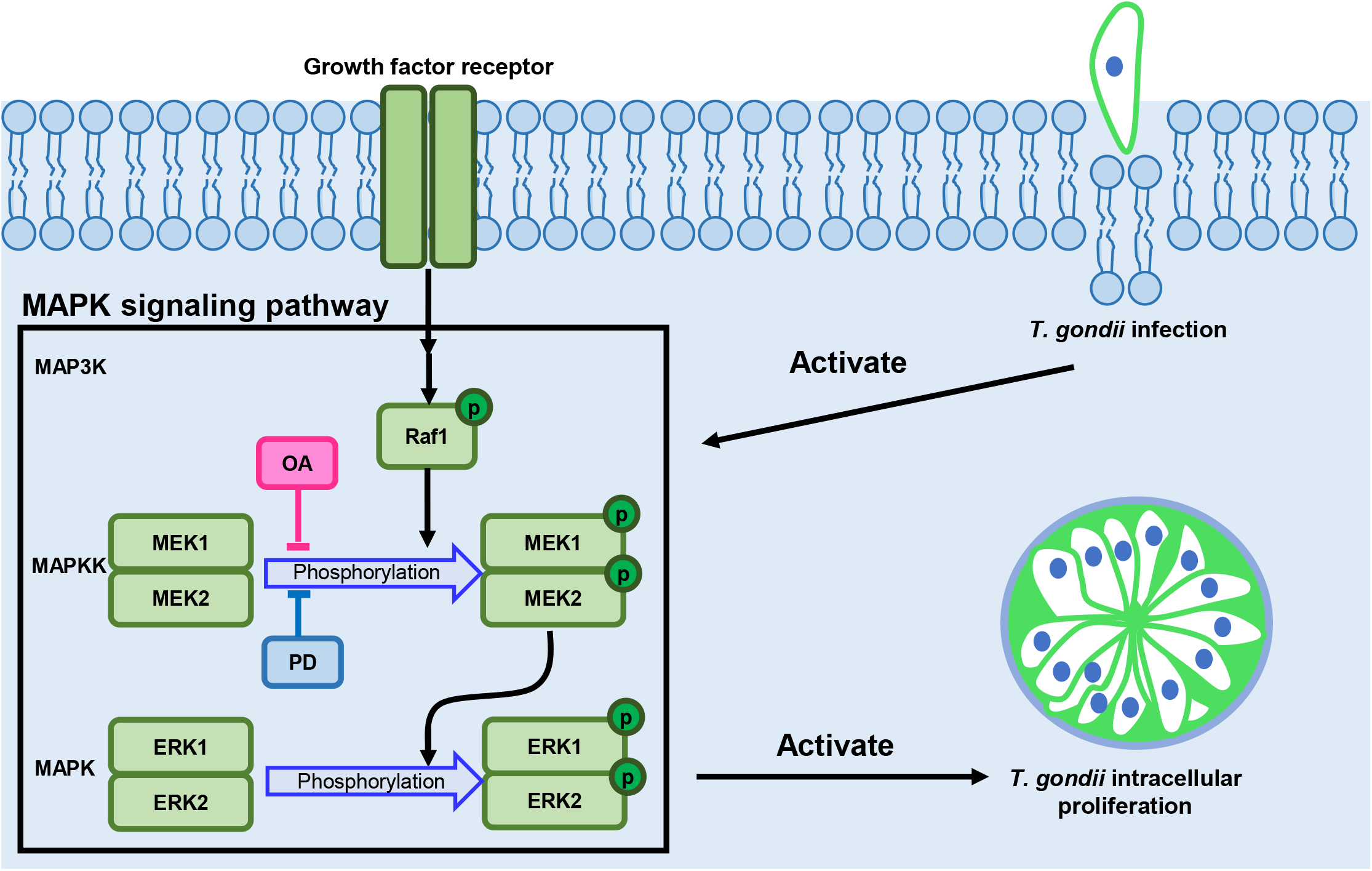
Schematic illustration model of the mode-of-action of OA on *T. gondii* proposed by this study. *T. gondii* infection activates the MAPK signaling pathway in the host cell. Phosphorylation of the host ERK1/2 promotes intracellular proliferation of *T. gondii*. Our findings suggest that OA inhibits the host ERK1/2 phosphorylation via binding to the allosteric site of host MEK1. This type of pathogen control by targeting host factors is known as host-directed therapy (HDT). This model proposed by this study shows the potential for the development of a new strategy, HDT, against *Toxoplasma*. PD: PD98059.

OA is a flavonoid compound known for its diverse pharmacological functions, including anticancer and anti-inflammatory properties^17^. In general, many flavonoids seem to cause protein kinase inhibition^32^. In addition, it has been reported that inhibition of the MAPK signaling pathway of host cells suppresses *T. gondii* proliferation^25^. Therefore, we hypothesized that OA inhibits the host MAPK signaling pathway, and inhibition of ERK phosphorylation by OA leads to the suppression of *T. gondii* proliferation. In fact, we demonstrated that OA inhibits the phosphorylation of host-cell ERK1/2 activated by *T. gondii* infection. Also, we focused on the structural similarity between OA and PD98059, a specific inhibitor of MEK1 possessing a flavone backbone. which provided further support for our hypothesis. In fact, we demonstrated that OA inhibits the phosphorylation of host-cell ERK1/2 activated by *T. gondii* infection, and its suppressive effect of OA on parasite proliferation remains unaffected by PD98059. Finally, *in silico* docking simulations showed that interacting energies of OA with MEK1 were comparable to those of PD98059. Taken together, the target molecule of OA is likely to be MEK1.

This study highlights the host-cell MAPK signaling pathway as a novel target for potential anti-*Toxoplasma* drug discovery. The MAPK signaling pathway demonstrates activation in human cancers, where MEK1/2 plays critical roles^33^. Therefore, inhibitors specifically targeting MEK1/2 have been developed, with trametinib earning approval from the US Food and Drug Administration (FDA) for clinical usage (FDA Reference ID: 4255758). The approval of trametinib demonstrates the feasibility of MEK1/2-targeted anti-toxoplasmosis drugs. However, the precise mechanisms underlying the inhibition of MAPK signaling pathway activation by *T. gondii* infection, resulting in the suppression of parasite proliferation, remain unclear. Our findings provide an impetus for further studies on the relationship between *T. gondii* infection and the host-cell MAPK signaling pathway, leading to drug development.

Recently, it has been established that intracellular pathogens, including viruses, bacteria, and parasites, rely on host cell factors for their proliferation^36^. Consequently, inhibitors targeting these essential host factors crucial for pathogens are considered potential drug candidates. This novel strategy, proposed as host-directed therapy (HDT), holds promise in reducing the emergence of drug resistance^37^. In parasitic diseases, this approach has been applied in a study focused on cutaneous leishmaniasis^38^, showing that tofacitinib, an inhibitor of the host cell Janus kinase 3, plays a critical role in the CD8 T cell IL-15 signaling pathway. This approach was found to be a safe strategy, effectively blocking immunopathologic responses locally while preserving protective responses. The mechanism of action identified for OA aligns precisely with this HDT strategy that has been underexplored for toxoplasmosis. Therefore, our findings could pave the way for HDT against *Toxoplasma*.

In conclusion, our findings revealed that suppressing host cell ERK phosphorylation effectively suppressed the intracellular growth of *Toxoplasma*. Given that OA does not affect the parasite directly, it is plausible that the emergence of drug resistance would be suppressed. Kinase inhibitors like OA and PD98059 hold potential as initial candidates to advance HDT studies in toxoplasmosis. Consequently, our study offers new insights into therapy against toxoplasmosis.

## MATERIALS AND METHODS

### Parasites

The archetypal type I strains of *T. gondii* RH and RH-GFP (ATCC #50940), along with the archetypal type II strain Fukaya, were cultured in Vero cells using high glucose Dulbecco’s Modified Eagle’s Medium (DMEM) (Sigma-Aldrich Co. St. Louis, MO). The medium contained 2% (v/v) fetal bovine serum (FBS) (Biowest, Nuaillé, France), 100 U/ml penicillin, and 100 µg/ml streptomycin (FUJIFILM Wako Pure Chemical Co., Osaka, Japan). All cells were maintained in a humidified incubator with 5% CO_2_ at 37°C. To harvest *Toxoplasma* tachyzoites, *T. gondii*-infected Vero cells were gently scraped ten times using a 25-gauge needle and a 10-ml syringe. The resulting cell solution was centrifuged at 150 × g for seven min to remove cell debris, and the supernatant, including tachyzoites, was collected. The concentration of tachyzoites was assessed using a hemocytometer, and the solution was diluted using DMEM with 2% (v/v) FBS.

Upon reaching confluency, *T. gondii* tachyzoites were placed onto Vero cells at a density of 10^5^/cm^2^ and then incubated with 5% CO_2_ at 37 °C. Following a 3-h incubation for parasite invasion, we performed a single wash of the plate using phosphate-buffered saline (PBS) (+) to eliminate any uninvaded tachyzoites. Subsequently, OA (MedChemExpress, NJ, CAS# 480-11-5) or PYR (Tokyo Chemical Industry, Tokyo, Japan) was added to each well. The culture medium, with or without these compounds, was replaced every two days.

### SRB cytotoxicity assay

Stock solutions of OA (50 mM) and PYR (2 mM) were prepared using DMSO and stored at –80 °C until needed. To assess the cytotoxicity of OA in host cells (Vero), we examined the SRB (Tokyo Chemical Industry) cytotoxicity assay^22,23^. Briefly, Vero cells were seeded at a density of 10^5^/cm^2^ in 24-well plates (Thermo Fisher Scientific, MA) and maintained in DMEM supplemented with 2% FBS. After two days of cell seeding, we confirmed the cell condition, i.e., confluence, before adding various concentrations (5 to 100 µM) of OA, PYR as an effective compound for anti-toxoplasmosis, or DMSO as a negative control to each well. These cells were cultured for six days. The SRB colorimetric assay was performed as described previously^23^.

### Monolayer disruption assay

Following the invasion of the *T. gondii* RH strain into Vero cells, as described in the “Parasites” subsection, we cultured the parasites in 12-well culture plates for six days. This culture was conducted with and without PYR (2 µM) or several concentrations of OA (5, 10, 25, 50, and 100 µM). Subsequently, the cultured cells were washed five times with 2 mL PBS (+) to eliminate unattached cells. The cells that remained attached to the plate were fixed by adding 1 ml of a 10% formalin neutral buffer solution (FUJIFILM Wako Pure Chemical Co.) and left to stand for 24 h at room temperature. The fixed cells were stained with a 1% aqueous solution of methylene blue (FUJIFILM Wako Pure Chemical Co.) for one hour. Images of the stained cells in each well were captured from the bottom side of the culture plate, and the coverage of stained cells in each well was analyzed using ImageJ software (National Institutes of Health, Bethesda, Maryland).

### Evaluation of the integrated density of *Toxoplasma* GFP signal and PV sizes in a host cell

The intracellular proliferation of the *T. gondii* RH-GFP strain was assessed in 12-well culture plates. Confluent Vero cell Parasites were invaded by the parasites and treated with or without PYR (2 µM) or different concentrations of OA (10 or 50 µM) for 48 h. Following this treatment, nonadhesive parasites, cells, and cellular debris were removed by washing twice with PBS (+). For each treatment group, images of five randomly selected fields were captured, and the average GFP signal integrated density was measured and calculated using ImageJ software. To determine percentages, the value of the GFP signal integrated density obtained from each compound-treated infection group was divided by the value obtained from the DMSO-treated infection group.

To analyze the PV sizes of *T. gondii* within individual host cells, Vero cells in 12-well culture plates were infected with RH-GFP parasites. These infected cells were treated with or without PYR (2 µM) or varying concentrations of OA (10 or 50 µM) for 30 h. The infected cells were washed twice using PBS (+) to remove nonadhesive cells and extracellular parasites. The cell nuclei were stained with a 0.05% Hoechst 33342 solution (Thermo Fisher Scientific) in PBS (+) for 5 min. In each treatment group, fluorescence images of ten randomly selected fields were captured, and the sizes of 100 PV were measured using ImageJ software.

### Evaluation of the effect of an ERK phosphorylation inhibitor on *Toxoplasma* proliferation

PD98059 was dissolved in DMSO to make a stock solution. The evaluation process of PD98059’s effect on *T. gondii* proliferation was identical to the previously described method but in a 6-well culture plate. The cells were harvested from culture plates by washing twice with ice-cold PBS (-) on ice and centrifuged at 4°C, 5,000 × g, for three min. Cell lysis was achieved by incubating the cell pellet in 120 μL of 1% Triton-X 100 buffer (1% Triton-X 100, 50 mM Tris-HCl (PH 7.5), 150 mM NaCl, 1 mM EDTA, 1 mM phenylmethylsulphonyl fluoride, 1 × protease inhibitor) on ice for 15 min, followed by three 10-second cycles of ultrasonic treatment. The total protein concentration was quantified using a bicinchoninic acid assay kit (Thermo Fisher Scientific). The protein samples were combined with 3 × SDS sample buffer containing 5% 2-mercaptoethanol and incubated at 95°C for 5 min. The protein was loaded onto an SDS-PAGE gel using 5% stacking and 8% running gel. The proteins in the gel were then transferred to a methanol-activated polyvinylidene difluoride membrane. The membrane was treated with blocking buffer (1% nonfat milk in PBS (-) with 0.1% Tween-20 solution (PBST)) for 10 min at room temperature. The membrane was incubated with the primary antibodies (diluted in blocking buffer) for a minimum of one hour at room temperature on a shaker. The following primary antibodies used were mouse monoclonal antibody ERK 1/2 (1:1,000 dilution, sc-514302; Santa Cruz Biotechnology, Inc., CA), mouse monoclonal antibody p-ERK 1/2 (1:1,000 dilution, sc-7383; Santa Cruz Biotechnology, Inc.), and rabbit polyclonal antibody beta-actin (1:5,000 dilution, 5057S; Cell Signaling, Inc., MA). After gently washing the membrane three times with PBST for 10 min, it was incubated with secondary antibodies (1:1,000 dilution in blocking buffer) for 30 min at room temperature. Goat anti–mouse IgG-horseradish peroxidase (HRP) (115-035-003; Jackson, Inc, PA) and goat anti–rabbit IgG-HRP (111-035-003; Jackson, Inc) were used as secondary antibodies. The membrane was then exposed to Immobilon Western Chemiluminescent HRP Substrate (Merck, Darmstadt, Germany) for one min, and the results were visualized using a luminescent image analyzer LAS4000 (FUJIFILM Wako Pure Chemical Co.).

### *In silico* docking simulation

The three-dimensional structure of the trametinib/MEK 1 complex was generated using Molecular Operating Environment (MOE), version 2022.02 (CCG Inc, Montreal, Canada), utilizing the Brookhaven Protein Databank 7JUR^28^ as a reference. Docking simulations within MOE were executed to mimic the trametinib-binding site of MEK 1 for PD98059 and OA. Following this, the ligand interaction mode within MOE was used to assess the physical and chemical parameters of trametinib, PD98059, and OA for MEK 1.

### Mice

Female C57BL/6J mice, weighing 20 ± 2 g and aged between 9 to 11 weeks, were obtained from SLC (Hamamatsu, Japan). These mice were kept in controlled temperature and humidity, under a 12-h day/night cycle, and provided with unlimited access to food and water. All studies were conducted according to protocols approved by Chiba University.

### Infection of *Toxoplasma* into mice

To prepare the cysts, mice were orally infected with cysts of the archetypal type II *T. gondii* strain Fukaya. Two months post-infection, the entire brain was extracted from the infected mice and homogenized in 10 mL PBS (-) to create a brain suspension. The cyst count was determined using a microscope. The suspension was then centrifuged at 440 × g for 5 min, diluted to a concentration of 20 cysts in 500 µL suspension, and orally administered to new mice. These newly infected mice were housed for four days prior to treatment.

### Compound treatment

To prepare the compound solution, a 6.5% DMSO in water was used as the solvent. Mice were orally administered with 500 µL of 6.5% DMSO as a negative control, 1 mg/kg/day of PYR, and 40 mg/kg/day of sulfadiazine in 6.5% DMSO as a positive control, or 50 mg/kg/day of OA in 6.5% DMSO orally once a day during 4-10 dpi. The body weight of the mice was measured daily until 12 dpi. The survival curves were plotted by monitoring the mouse health condition until 30 dpi. All animal treatments adhered to the guidelines established set forth by the Chiba University Animal Ethics Committee.

### Statistical analyses

Statistical analyses were conducted using OriginPro, Version 2021 (OriginLab Corporation, MA). All data except survival rate were analyzed using the ANOVA test. Survival analysis was performed using the Kaplan–Meier method, and the log-rank test was used for making comparisons. Data with *P* < 0.05 were considered statistically significant.

## AUTHOR INFORMATION

### Corresponding Author

**Kenji Hikosaka**

Department of Infection and Host Defense, Graduate School of Medicine, Chiba University, 1-8-1 Inohana, Chuo-ku, Chiba 260-8670, Japan.

### Authors

**Ziyue Z Zhang**

Department of Infection and Host Defense, Graduate School of Medicine, Chiba University, 1-8-1 Inohana, Chuo-ku, Chiba 260-8670, Japan

**Kazumi Norose**

Department of Infection and Host Defense, Graduate School of Medicine, Chiba University, 1-8-1 Inohana, Chuo-ku, Chiba 260-8670, Japan; Department of Parasitology, Shinshu University School of Medicine, 3-1-1 Asahi, Matsumoto 390-8621, Japan

**Noriko Shinjyo**

Laboratory of Immune Homeostasis, WPI Immunology Frontier Research Center, Osaka University, Osaka 565-0871, Japan; School of Tropical Medicine and Global Health, Nagasaki University, Nagasaki 852-8523, Japan; https://orcid.org/0000-0003-4501-4513

**Xiaoxia X Lin**

Department of Infection and Host Defense, Graduate School of Medicine, Chiba University, 1-8-1 Inohana, Chuo-ku, Chiba 260-8670, Japan

**Akiko Suganami**

Department of Bioinformatics, Graduate School of Medicine, Chiba University, 1-8-1 Inohana, Chuo-ku, Chiba 260-8670, Japan

**Yutaka Tamura**

Department of Bioinformatics, Graduate School of Medicine, Chiba University, 1-8-1 Inohana, Chuo-ku, Chiba 260-8670, Japan; https://orcid.org/0000-0001-5373-6909

**Hirokazu Sakamoto**

Department of Infection and Host Defense, Graduate School of Medicine, Chiba University, 1-8-1 Inohana, Chuo-ku, Chiba 260-8670, Japan; Department of Pathology, Stanford School of Medicine, Stanford, California 94305, USA; https://orcid.org/0000-0001-9368-105X

### Author Contributions

Z.Z.Z. and K.H. conceived the concept of this study. Z.Z.Z., K.N., N.S., X.X.L., H.S., and

K.H. designed the experiments. A.S. and Y.T. conducted the *in silico* docking simulation analyses. Z.Z.Z. prepared first draft of the manuscript and the Figures. All authors edited the manuscript and have approved the summitted manuscript.

### Notes

The authors declare no competing financial interest.

## ABBREVIATIONS

Am: *Astragalus membranaceus*
FBS: fetal bovine serum
HDT: host-directed therapy
MEK: mitogen-activated protein kinase/extracellular signal-regulated kinase
MOE: Molecular Operating Environment
OA: oroxylin A
PV: parasitophorous vacuole
PYR: pyrimethamine
Sb: *Scutellaria baicalensis*
TCM: Traditional Chinese medicine
Tg/*T. gondii*: *Toxoplasma gondii*

## ACKNOWLEDGEMENTS

This work was supported by JSPS KAKENHI (Grant Number JP19K07520, JP19K16627, JP21KK0285, JP22H04633), AMED (Grant Number JP21fk0108569 and JP22jm0610076), Ohyama Health Foundation, Hirose Foundation, Future Medicine Education and Research Organization of Chiba University.

