## Supplementary materials for "Suppressive effects of oroxylin A on intracellular proliferation of *Toxoplasma gondii* via host cell ERK phosphorylation inhibition"

**Methods**

**Supplementary 1. Pretreatment of compounds before and while *Toxoplasma* invasion**

*T. gondii*-infected Vero cells were scrapped ten times rapidly using a 25-gauge needle with a 10 mL syringe. The cell solution was centrifuged at 150 × g for seven min to remove cell debris, and the supernatant including tachyzoites was collected. The supernatant was centrifuged at 1,000 × g for five min to collect *T. gondii* tachyzoites. The pretreatment (PT) was performed by re-suspending *T. gondii* pellet into DMSO, OA (50 μM), or PYR (2 μM) added medium to make a 4 × 10^5^/mL suspension and these *T. gondii*-suspended solutions were incubated at 37^o^C for 1 h.

For the compound un-treated culture while invasion (UT), the tachyzoites were centrifuged, dissolved in DMEM, and added to confluent Vero cells in 24-well plates. After 3 h incubation for parasite invasion, we washed the plate twice using PBS (+) to remove uninvaded tachyzoites. Infected cells were then incubated for 48 h using a DMEM until staining and imaging were performed.

The compound treated culture while invasion (T) was performed by adding compound pretreated suspension into confluent Vero cells in 24-well plates, and infected cells were washed twice using PBS (+) to remove uninvaded *T. gondii*. Infected cells were then incubated for 48 h using DMEM until staining and imaging were performed.

**Supplementary 2. Cytotoxicity of OA evaluated by SRB cytotoxicity assay**

Stock solutions of OA (50 mM), PYR (2 mM), and PD98059 (50 mM) were prepared using DMSO and stored at -80 ºC until use. Cytotoxicity of OA on host cells (Vero) was examined using the SRB cytotoxicity assay^1^. Briefly, Vero cells were seeded at 10^5^/cm^2^ to 24-well plates and cultured in DMEM with 2% FBS. After two days of seeding cells, we confirmed the cell condition, that is, confluent, and added several concentrations of OA, PD98059, and PYR as an effective compound for toxoplasmosis, or DMSO as a negative control to each well. These cells were cultured for six days. SRB colorimetric assay was performed as described previously^1^.

**Supplementary 3. The p-ERK 1/2 level change in *T. gondii*-infected HFF cells treated with different compounds**

Evaluation of the effect of PD98059 on *T. gondii* proliferation was performed in a 6-well culture plate. HFF cells were seeded and cultured for two days until confluence, 10^5^/cm^2^ *T. gondii* RH-GFP strain were then seeded and allowed to invade for three h and washed the plate twice using PBS (+) to remove uninvaded tachyzoites. The compound treatment was performed for 48 h.

HFF cells were isolated from culture plates by washing twice using ice-cold PBS (-) on ice and centrifuged at 4 ^o^C, 5,000 × g, for three min. Lysis was performed by incubating the cell pellet in 120 μL 1% Triton-X 100 buffer (1% Triton-X 100, 50 mM Tris-HCL (PH 7.5), 150 mM NaCl, 1 mM EDTA, 1 mM phenylmethylsulphonyl fluoride, 1 × protease inhibitor) on ice for 15 min, and ultrasonic treatment was performed 10 s three times during treatment. The whole protein concentration was quantified using a bicinchoninic acid assay kit according to the manual. Protein samples were then mixed with 3 X SDS sample buffer with 5% 2-mercaptoethanol and incubated at 95 ^o^C for five min. The protein was loaded and ran SDS-PAGE using 5% stacking and 8% running gel. After that, the protein in the gel was transferred onto a methanol-activated polyvinylidene difluoride membrane. The membrane was treated with blocking buffer (1% non-fat milk in PBS with 0.1% Tween-20 solution (PBST)) for 10 min at room temperature. The membrane was incubated with primary antibody (dilution in blocking buffer) for at least one h at room temperature on a shaker. The following primary antibodies were used: mouse monoclonal antibody ERK 1/2 1:1,000, mouse monoclonal antibody p-ERK 1/2 1:1,000, and rabbit polyclonal antibody beta-actin 1:5,000. Then, the membrane was washed gently using PBST for 10 min three times and then incubated with a secondary antibody (1:1,000 dilution in blocking buffer) for 30 min at room temperature. Goat anti-mouse IgG-HRP and goat anti-rabbit IgG-HRP were used as secondary antibodies. The membrane was subsequently incubated with Immobilon Western Chemiluminescent HRP Substrate for one min, and the result was detected by a luminescent image analyzer LAS4000.

**Supplementary 4. Monolayer disruption assay of OA and PD98059 using Fukaya strain**

After invasion of the *T. gondii* Fukaya strain using 12-well culture plates, we cultured parasites for six days with or without PYR (2 µM) or several concentrations of OA (5, 10, 25, 50, and 100 µM). Cultured cells were washed five times using 2 mL PBS (+) to remove unattached cells. Subsequently, cells attaching to the plate were fixed by adding 1 ml 10% formalin neutral buffer solution (Wako, Osaka, Japan) and stood for 24 h at room temperature. Fixed cells were stained using methylene blue 1% aqueous solution (Wako) for one hr. Images of stained cells in each well were taken from the bottom side of the culture plate, and the bottom coverage of stained cells on each well was analyzed using the software ImageJ.

**Legends of supplementary figures**

**Fig. 1** No significant changes between DMSO, OA, and PYR pretreatment before or while invasion. PT. pretreated group. UT, pretreated before invasion but un-treated while invasion group. T, pretreated before invasion and treated while invasion group. *T. gondii* RH-GFP was used in this experiment, and proliferation was estimated by GFP signal. Scale bar, 100 µm.

**Fig. 2** No cytotoxicity was observed except 100 µM OA treatment and 50 µM OA + 25 µM PD98059 cotreatment groups. Experiments were repeated three times independently and triplicated in each experiment with similar results., and the representative data is shown. Data are presented as mean ± SEM. The white bar indicates a result of no compound supplementation as a negative control, and the black bar indicates a result of PYR supplementation. *****, *P <* 0.001 vs untreated group.

**Fig. 3** OA downregulated ERK 1/2 phosphorylation in HFF cells as the host cell. All treatments were performed for 48 h. Cells in each well were lysed using 0.1 % triton buffer and concentration was checked by a bicinchoninic acid assay. After that western blotting was performed by adding 1.5 µg whole protein in each lane.

**Fig. 4** Monolayer disruption assay showed dose-dependent suppressive effects of OA on proliferation of *T. gondii* Fukaya strain. (A) Images of monolayer disruption assay. Experiments were repeated independently two times and triplicated in each experiment, and the representative data is shown. (B) The % of the bottom coverages. Data are presented as mean ± SD.
