## Supplementary figures for "Suppressive effects of oroxylin A on intracellular proliferation of *Toxoplasma gondii* via host cell ERK phosphorylation inhibition"

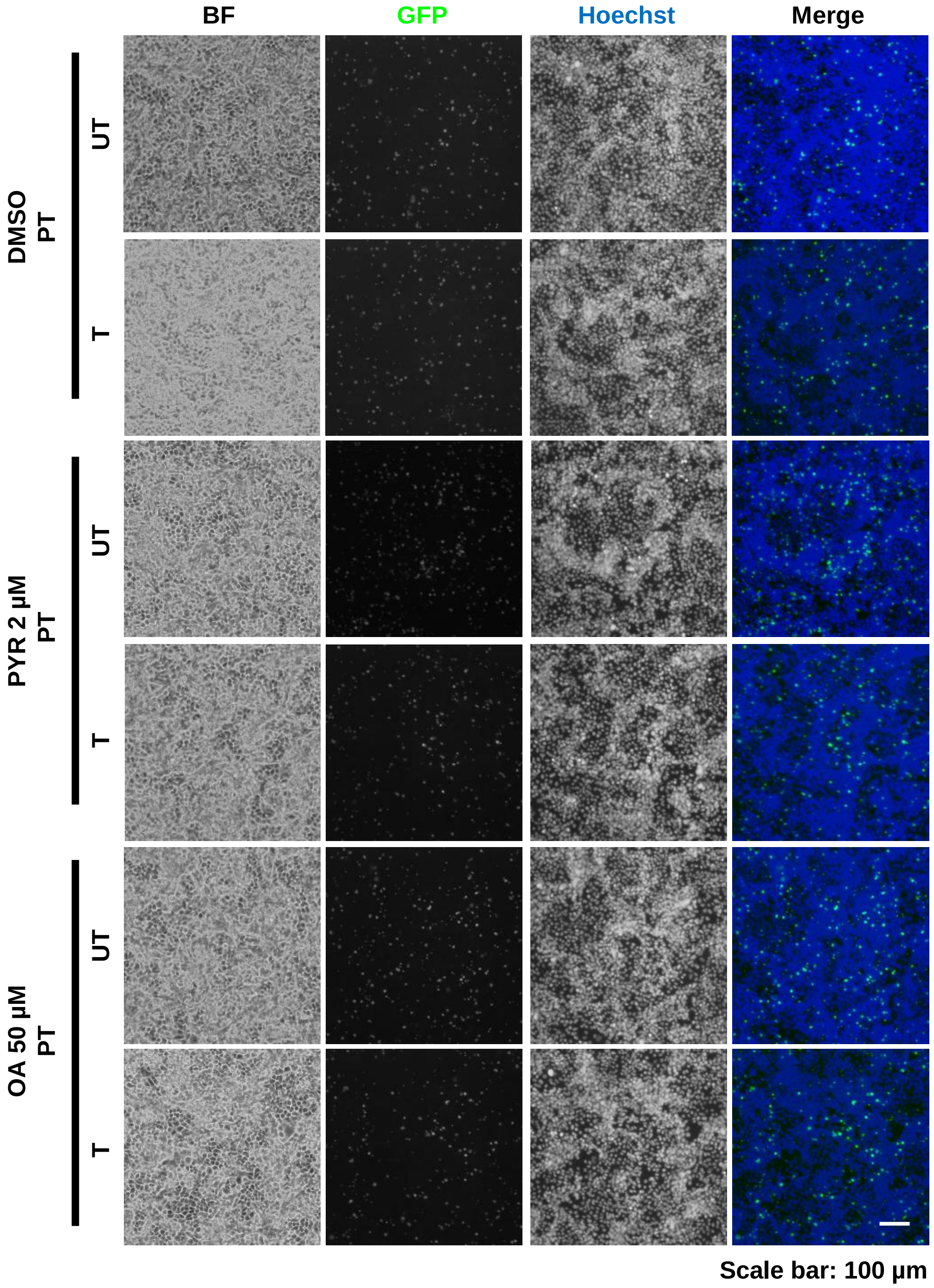


**Fig. 1** No significant changes between DMSO, OA, and PYR pretreatment before or while invasion. PT. pretreated group. UT, pretreated before invasion but un-treated while invasion group. T, pretreated before invasion and treated while invasion group. *T. gondii* RH-GFP was used in this experiment, and proliferation was estimated by GFP signal. Scale bar, 100 μm.

Supplementary Fig. 1_Zhang et al.

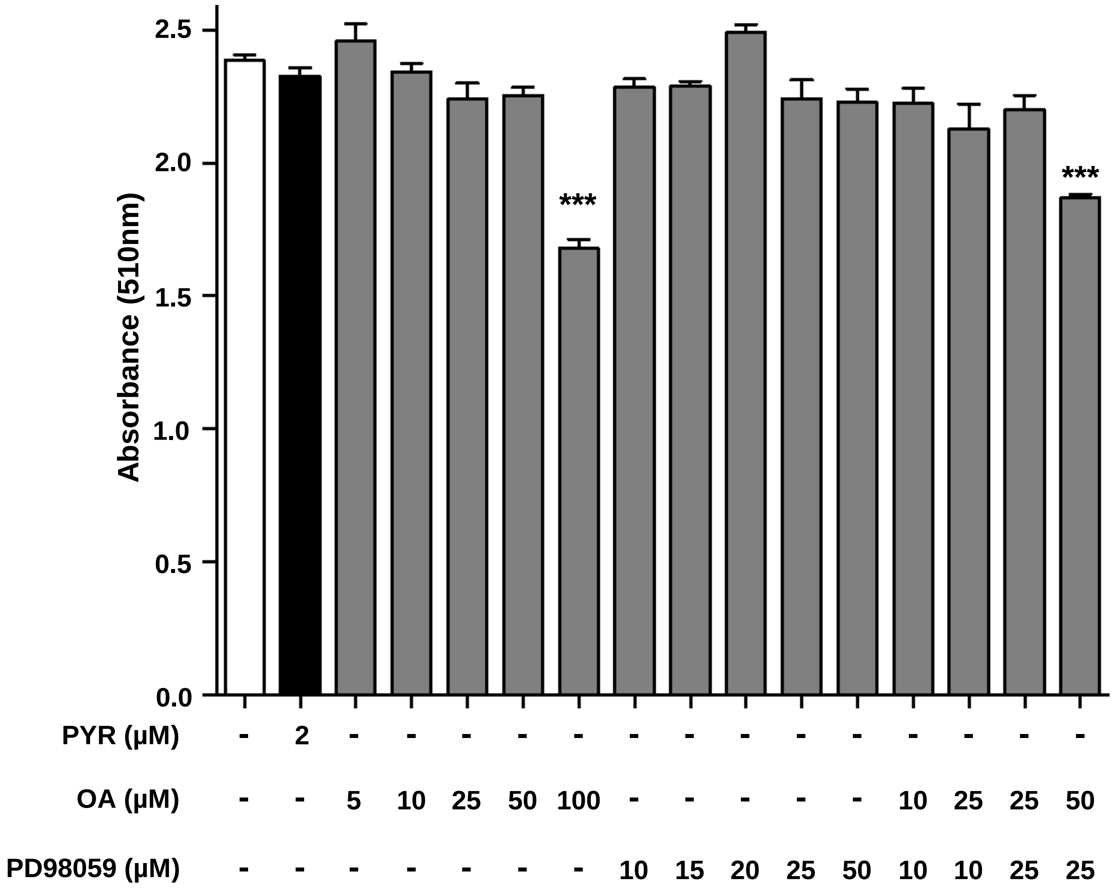


**Fig. 2** No cytotoxicity was observed except 100 μM OA treatment and 50 μM OA + 25 μM PD98059 cotreatment groups. Experiments were repeated three times independently and triplicated in each experiment with similar results., and the representative data is shown. Data are presented as mean ± SEM. The white bar indicates a result of no compound supplementation as a negative control, and the black bar indicates a result of PYR supplementation. ***, $P < 0.001$ vs untreated group.

Supplementary Fig. 2_Zhang et al.

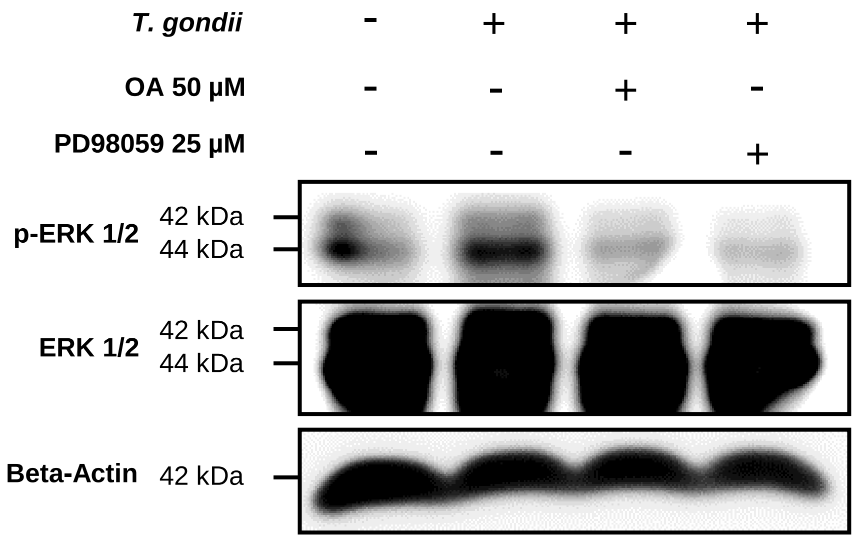


**Fig. 3** OA downregulated ERK 1/2 phosphorylation in HFF cells as the host cell. All treatments were performed for 48 h. Cells in each well were lysed using 0.1 % triton buffer and concentration was checked by a bicinchoninic acid assay. After that western blotting was performed by adding 1.5 µg whole protein in each lane.

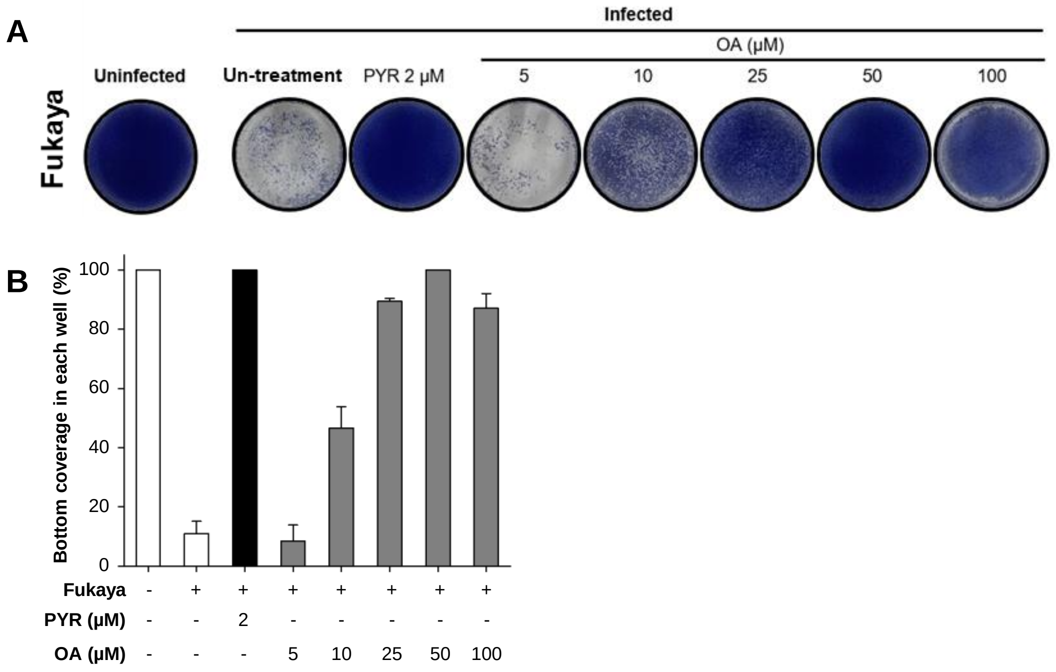


**Fig. 4** Monolayer disruption assay showed dose-dependent suppressive effects of OA on proliferation of T. gondii Fukaya strain. (A) Images of monolayer disruption assay. Experiments were repeated independently two times and triplicated in each experiment, and the representative data is shown. (B) The % of the bottom coverages. Data are presented as mean ± SD.

Supplementary Fig. 4_Zhang et al.
